# Characterizing the role of RSD-6 in the biogenesis of virus-derived small interfering RNAs and the modulation of viral pathogenesis

**DOI:** 10.1101/2025.09.02.673749

**Authors:** Krishna Dahal, Mingli Xia, Jinfeng Lu, Teng Yan, Rui Lu

## Abstract

Small interfering RNAs (siRNAs) produced through the processing of viral double-stranded RNAs mediate potent antiviral RNA interference (RNAi) in eukaryotes. In *Caenorhabditis elegans*, such an antiviral defense is further amplified through the production of secondary siRNAs, yet the mechanisms by which secondary virus-derived siRNAs (vsiRNAs) confer protection remain poorly understood. Here, we characterize the role of *rsd-6*, which encodes a Tudor domain protein and plays important role in antiviral RNAi, in vsiRNA biogenesis and modulation of viral pathogenesis. Using CRISPR Cas9-generated *rsd-6* null mutants, we show that both primary and secondary vsiRNAs accumulate normally in the absence of RSD-6, indicating that it functions downstream of secondary vsiRNA biogenesis. We further showed that secondary vsiRNAs generated in *rrf-1*-independent manner remained detected in the absence of RSD-6 and viral replication is further enhanced in *rrf-1;rsd-6* double mutants compared to *rrf-1* single mutants, suggesting a role of *rsd-6* in mediating antiviral guided by all secondary vsiRNAs. Consistently, *rsd-6* mutants exhibited more severe pathogenesis upon Orsay virus infection compared to *rrf-1* mutants, underscoring its role as a major determinant of viral disease outcome. Domain characterization established that the N-terminal tandem domains of RSD-6 are required for antiviral activity, while the C-terminal Tudor domains are dispensable. Functional conservation was confirmed in *C. briggsae*, where silencing of the *rsd-6* homolog enhanced viral replication. Together, our findings identify RSD-6 as a key effector acting downstream of secondary vsiRNA production and highlight its conserved role in modulating viral replication and pathogenesis across *Caenorhabditis* species.

**Importance:** In *C. elegans*, the RNAi-mediated antiviral defense relies on the production of secondary virus-derived siRNAs (vsiRNAs) to achieve an amplified antiviral effect. However, the mechanism by which these secondary vsiRNAs confer protection remains poorly understood. This is primarily due to the limited number of identified key effector genes. To address this knowledge gap, we profiled vsiRNA biogenesis in loss-of-function mutants and discovered that *rsd-6* is dispensable for the production of secondary vsiRNAs, suggesting a role of *rsd-6* in mediating antiviral defense downstream of secondary vsiRNA biogenesis. Worm survival assay further confirmed that *rsd-6* is a critical modulator of viral pathogenesis and its antiviral function is conserved across Caenorhabditis species. The RSD-6 protein features three N-terminal tandem domains of unknow function and two tandem Tudor domains at its C-terminus. Our domain analyses demonstrated that the N-terminal tandem domains, but not the C-terminal Tudor domains, are essential for antiviral function. The identification of *rsd-6* as a key effector gene acting downstream of vsiRNA biogenesis provides a solid foundation for elucidating the mechanism of antiviral RNAi amplification.

## Introduction

Small interfering RNAs (siRNAs) derived from replicating viruses mediate potent antiviral silencing across diverse organisms, including fungi, plants, invertebrates, and vertebrates (1–4). This siRNA-mediated antiviral immunity, collectively referred to as antiviral RNA interference (RNAi), is initiated when viral double-stranded RNA (dsRNA), often generated as replication intermediates, is processed by Dicer proteins into primary siRNAs, typically 21 to 24 nucleotides (nt) in length (4). These primary siRNAs are then loaded into Argonaute (AGO) proteins within the RNA-induced silencing complex (RISC), where they direct the cleavage of complementary viral RNA transcripts. In plants, RNA-dependent RNA polymerases (RdRPs) convert the cleaved RNA fragments into new dsRNAs, which are subsequently processed into secondary siRNAs (5, 6). In this way, antiviral RNAi is amplified.

Although key factors such as Dicer, AGO, RdRPs and dsRNA binding proteins (DRBPs) are conserved in nematode antiviral RNAi, they employ distinct mechanisms for the production of primary and secondary virus-derived siRNAs (vsiRNAs). In *C. elegans*, Dicer is required for the production of primary, but not secondary, vsiRNAs. Primary vsiRNA biogenesis also depends on DRH-1 (Dicer-related RNA helicase 1) and RDE-4 (RNAi defective 4), with DRH-1 playing a predominant role (7–9). These primary vsiRNAs are loaded into RDE-1, a worm AGO protein, and guide the cleavage of viral target transcripts by the endonuclease RDE-8 (10). RDE-3 also plays a critical role in antiviral defense, and recent studies suggest that it modifies cleaved viral transcripts by appending poly(UG) tails to their 3′ ends (11, 12). These poly(UG) tails subsequently recruit RRF-1, an RdRP, to synthesize secondary siRNAs in a Dicer-independent manner (13). Unlike primary siRNAs, which possess a single 5′ monophosphate, secondary siRNAs are uniformly 22 nucleotides long and carry a 5′ triphosphorylated guanosine, hence they are commonly referred to as 22G siRNAs (14, 15).

To date, four closely related RNA viruses have been identified as natural pathogens of nematodes (16–18). Among them, Orsay virus (OrV) is the first and best-characterized nematode virus that infects the intestinal cells of *C. elegans*. The other three viruses, Santeuil virus (SANTV), Le Blanc virus, and Melnik virus, infect *C. briggsae* and share similar intestinal tropism and genome organization. OrV is a small, non-enveloped icosahedral virus with a bipartite, positive-sense RNA genome. The RNA1 segment encodes an RNA-dependent RNA polymerase (RdRp) required for viral replication, while the RNA2 segment encodes both a capsid protein and a fusion protein essential for infection (19). Structurally and genetically, OrV resembles nodaviruses but displays unique features, including strict host specificity and the absence of subgenomic RNAs. The intestinal tropism of OrV is likely determined by factors required for viral genome replication rather than cell entry. This is supported by the observation that infection initiated from a transgene, bypassing the cell entry step, still remains restricted to the intestine (20).

In plants and insects, successful viral infection often depends on virus-encoded RNAi suppressors, which are also responsible for the diseases associated with infection (21, 22). In contrast, Orsay virus infects wild-type *C. elegans* at low efficiency and does not induce overt pathology (14). However, enhanced Orsay viral replication and disease symptoms were observed in the presence of exogenous RNAi suppressors or in mutants defective in RNAi (7, 14). This suggests that, unlike many plant and insect viruses, Orsay virus lacks a potent RNAi suppression activity. Notably, both OrV and SANTV were originally discovered in wild isolates of nematodes that are naturally susceptible to infection and display associated intestinal pathology (9, 16, 17). This demonstrates that viral pathogenesis can be studied in nematodes, although a viral counter-defense mechanism, e.g., an RNAi suppressor, has yet to be identified.

Owing to its high efficiency in gene discovery through genome-wide genetic screens, the *C. elegans* system has yielded the most identified antiviral RNAi genes of any model system (23). Functional and mechanistic studies of these genes have greatly advanced our understanding of the biogenesis and activity of primary vsiRNAs. In contrast, while much has been learned about how secondary vsiRNAs are produced, how they mediate antiviral silencing remains largely unknown. This knowledge gap is partly due to the limited number of identified *C. elegans* genes that function downstream of secondary vsiRNA biogenesis. Here, we report the characterization of *C. elegans rsd-6* in vsiRNA biogenesis and its role in modulating viral pathogenesis. Our findings suggest that *rsd-6*, as a conserved antiviral factor in *Caenorhabditis* nematodes, functions downstream of secondary vsiRNA biogenesis and serves as a key modulator of viral pathogenesis.

## Results

### Generation of loss of function allele for rsd-6

We previously demonstrated that premature stop codons resulting from single nucleotide changes render *rsd-6* non-functional in antiviral defense (12). To facilitate reliable genotyping of *rsd-6* loss-of-function mutants, we sought to generate a defined *rsd-6* null allele using CRISPR– Cas9 genome editing. For this purpose, we designed CRISPR RNAs (crRNAs) that would not only delete essential coding sequences but also introduce a downstream frameshift mutation (Figure 1A). A mixture of crRNAs, tracrRNA, single-stranded DNA donor, Cas9 protein, and PRF4 plasmid was injected into wild-type N2 worms carrying the FR1gfp replicon transgene (14). F1 progeny exhibiting the roller phenotype indicated successful formation of extrachromosomal transgene arrays.

**Figure 1.**
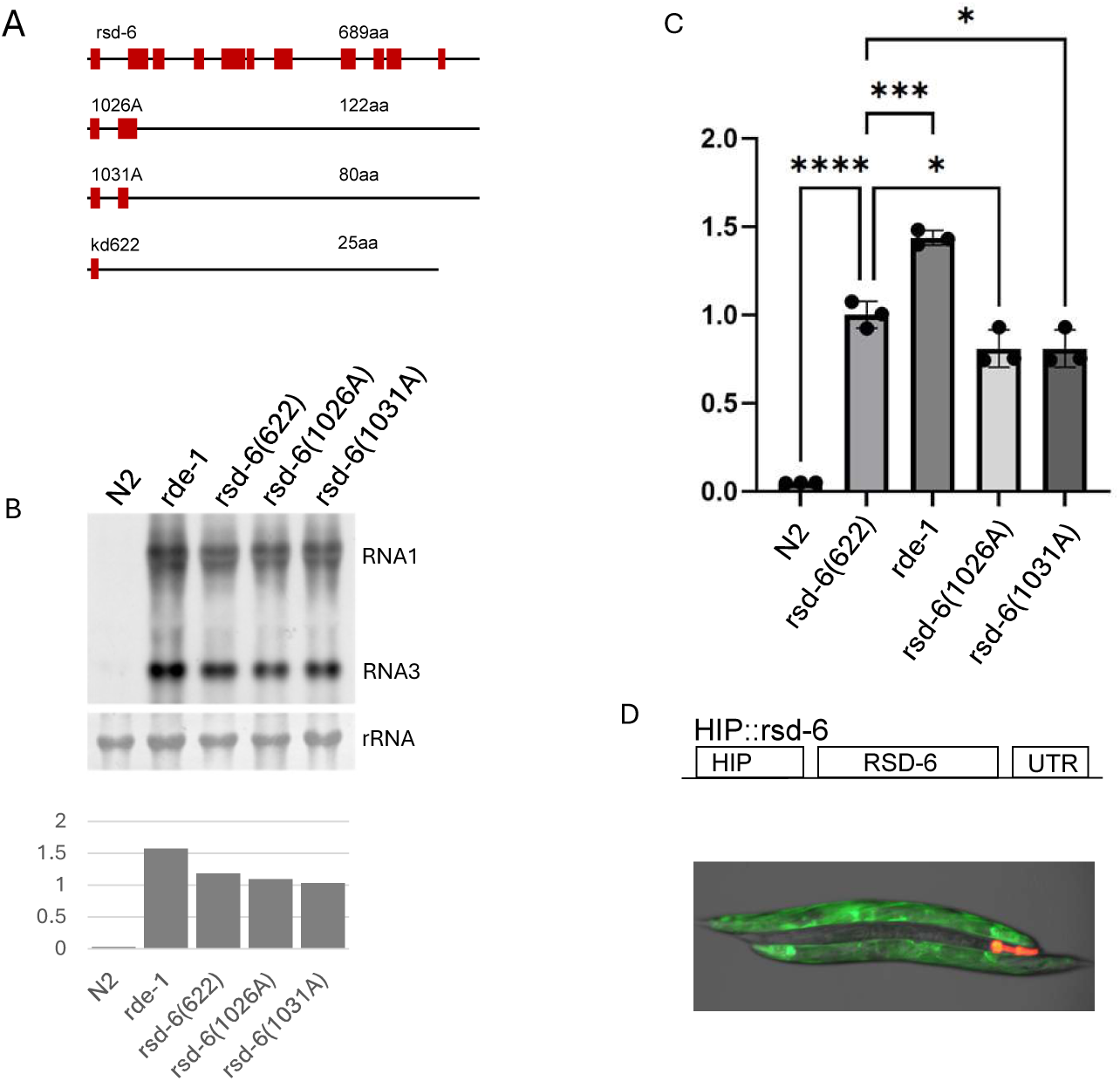
Generation and characterization of an rsd-6 null allele using genome editing. A. Schematic representation of the wild-type and mutant alleles of the *rsd-6* gene. Exons are shown as solid boxes; introns and untranslated regions are represented by lines. The predicted protein sizes from conceptual translation are indicated for each allele. B. Upper panel: Northern blot detection of FR1gfp genomic and subgenomic RNAs in wild-type N2 and *rsd-6* mutant (*kd622)*) strains. A labeled cDNA probe derived from the GFP coding sequence was used.in this northern blotting assay. Lower panel: Quantification of FR1gfp subgenomic RNA3 levels from the northern blot using ImageJ. C. RT-qPCR quantification of FR1gfp genomic RNA1 accumulation in wild-type and *rsd-6(kd622)* mutants. A primer pair specific for FR1gfp RNA1 was used. Amplification of the endogenous *act-1* gene served as an internal control. ****, P < 0.00001.***, *p < 0.0001.* *, P < 0.01. D. Upper panel: Schematic of the *rsd-6* transgene under control of a heat-inducible promoter (HIP). UTR: 3′ untranslated region of *unc-54*. Lower panel: Overexpression of wildtype *rsd-6* rescues antiviral RNAi in *rsd-6(kd622)* mutants (with red fluorescence in head). The image was recorded 24 hours post heat induction.

To identify worms carrying the intended deletion, we performed PCR genotyping using primers flanking the targeted region. Sanger sequencing confirmed the deletion in *rsd-6* genomic DNA. We next assessed FR1gfp replication by northern blotting and RT-qPCR. As shown in Figures 1B and 1C, FR1gfp replication was markedly elevated in both *rde-1*(*ok569*) null mutants and in previously characterized *rsd-6* premature stop codon mutants (12), compared to wild-type N2 animals. Importantly, a similar increase was observed in animals carrying the newly generated *rsd-6* deletion allele, designated *kd622*. To confirm that the enhanced FR1gfp replication in *kd622* animals was due specifically to loss of *rsd-6* function, we introduced a transgene array expressing wild-type *rsd-6* under a heat-inducible promoter (HIP) by gonad microinjection (Figure 1D, upper panel). This array also carried an mCherry marker expressed in the head region. Following heat induction, *kd622* animals harboring the *HIP::rsd-6* transgene exhibited no detectable FR1gfp replication (Figure 1D, lower panel), confirming that the antiviral defect in *kd622* animals is indeed attributable to loss of *rsd-6*. The *kd622* allele contains a 1,506 bp deletion that removes all coding sequences from the second and third exons and parts of the first and fourth exons. This deletion is predicted to also introduce a frameshift mutation, resulting in a severely truncated polypeptide of only 25 amino acids, if translated. Together, these findings establish *kd622* as a bona fide *rsd-6* null allele.

### vsiRNAs derived from Orsay virus RNA1 and RNA2 exhibit distinct size distributions and abundances

Currently, the precise role of *rsd-6* in antiviral RNAi remains unclear. To test whether *rsd-6* contributes to the biogenesis of viral siRNAs (vsiRNAs) during natural infection, we analyzed the abundance of Orsay virus-derived siRNAs through deep sequencing. Small RNAs, including miRNAs and vsiRNAs, were enriched from total RNA prepared from various *C. elegans* strains using a standardized precipitation protocol (7). The enriched small RNAs were treated with polyphosphatase prior to sequencing, a step that enables capture of secondary vsiRNAs, which carry a 5′ triphosphate that otherwise prevents adapter ligation. Each small RNA library contained between 9 and 41 million reads (Table S1), ensuring comprehensive coverage of vsiRNAs.

Sequencing data were analyzed with an in-house developed bioinformatics pipeline to identify both primary and secondary siRNAs derived from the Orsay virus RNA1 and RNA2 segments. In both wild-type N2 animals and RNAi-defective mutants, we observed that RNA1 generated significantly fewer primary vsiRNAs (22–24 nt) than RNA2 (Figure S1), indicating that production of RNA1-derived primary vsiRNAs is comparatively inefficient. In contrast, RNA1 consistently yielded a much higher level of minus-strand 22G vsiRNAs, predominantly secondary siRNAs, relative to RNA2, across all strains except *rde-1* and *rrf-1* mutants, in which secondary vsiRNAs were nearly absent. This finding is consistent with prior studies showing that RDE-1, together with RDE-8 and RDE-3, acts upstream of RRF-1 to generate templates for secondary siRNA production (10, 11, 23). Taken together, these data demonstrate that, relative to its length, Orsay virus RNA1 (3,421 nt) produces fewer primary vsiRNAs but substantially more secondary vsiRNAs than RNA2 (2,574 nt), revealing segment-specific differences in vsiRNA biogenesis during RNAi-mediated antiviral defense.

### RSD-6 is dispensable for the biogenesis of primary and secondary vsiRNAs

The presence of both primary and secondary vsiRNAs in *rsd-6* (*kd622*) mutants (Figure S1) prompted us to test whether *rsd-6* functions downstream of secondary vsiRNA biogenesis in antiviral RNAi. To address this, we performed quantitative analyses of OrV RNA1-derived vsiRNAs in several RNAi-defective mutants, including *rsd-6* (*kd622*) animals. We focused on OrV RNA1 because RNA2 generates relatively few secondary vsiRNAs, complicating quantitative interpretation (Figure S1). For cross-sample comparisons, vsiRNA read counts were normalized to one million endogenous *C. elegans* miRNAs, whose expression is unaffected by the mutations analyzed.

Primary vsiRNAs, which carry a 5′ monophosphate, were cloned and sequenced without enzymatic pretreatment. Despite differences in OrV replication levels among wild-type N2, *rde-1*, *rrf-1*, and *rsd-6* mutants (Figure 2A), the size distribution of primary vsiRNAs was consistent across all strains (Figure 2B). Quantification of 22–24 nt primary vsiRNAs from OrV RNA1 revealed that *rsd-6* mutants accumulated these molecules at levels comparable to *rde-1* and *rrf-1* mutants (Figure 2D, left panel). Because *rde-1* and *rrf-1* are dispensable for primary vsiRNA production (9, 14, 24), these results suggest that *rsd-6* is likewise not required for primary vsiRNA biogenesis. Furthermore, primary vsiRNAs from all genetic backgrounds mapped to similar regions of the OrV RNA1 and RNA2 genomes, displaying unique patterns of genome coverage (Figure S2 and data not shown). Even wild-type N2 animals, which produced fewer primary vsiRNAs overall, exhibited the same mapping profiles. This finding indicates that the sequence features of viral double-stranded RNA, rather than its abundance, determine which genomic regions contribute to the production of primary vsiRNAs.

**Figure 2.**
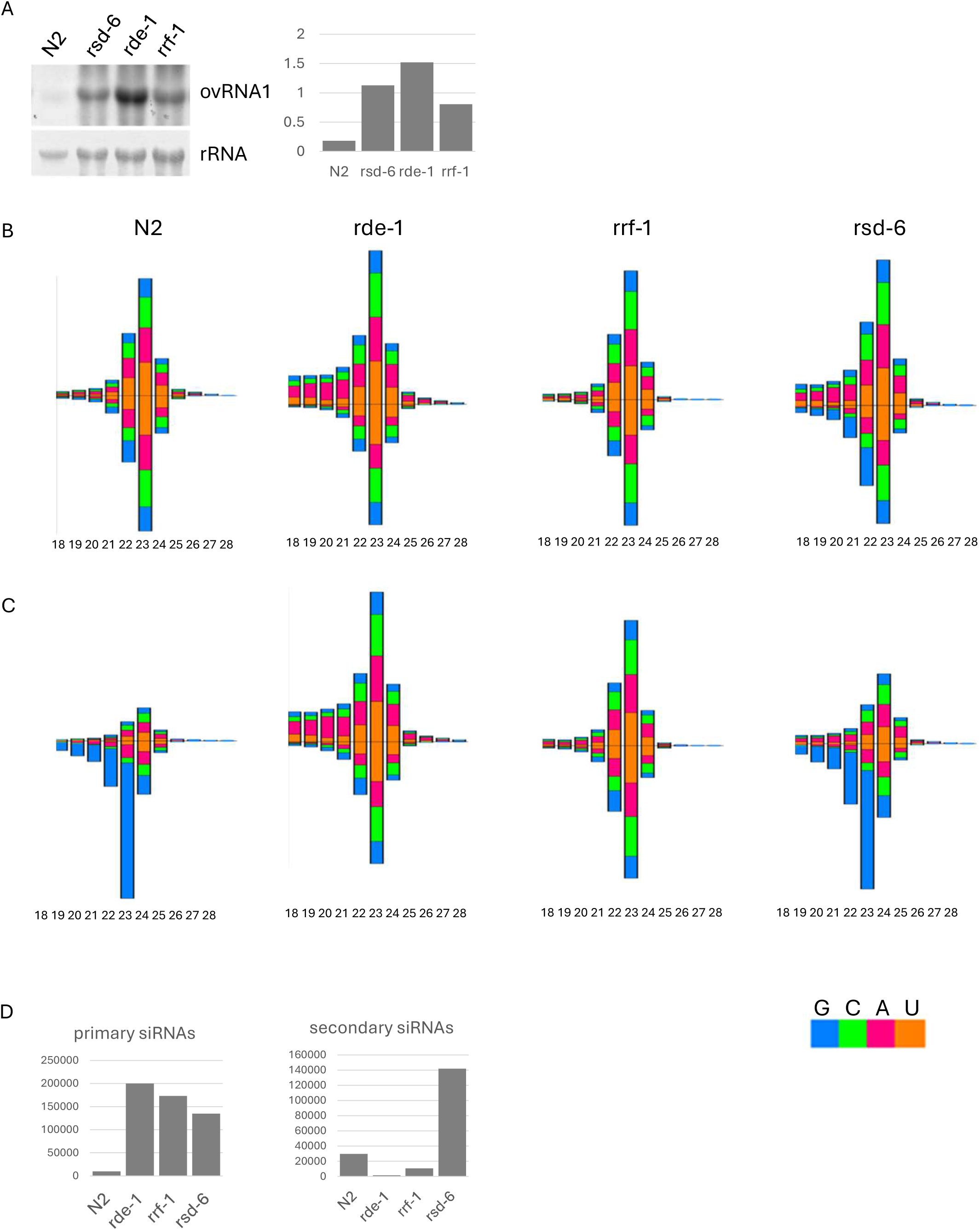
Quantification of siRNAs derived from Orsay virus genomic RNA1 in different genetic backgrounds. A. Left panel: Northern blot analysis of Orsay virus genomic RNA1 in the indicated *C. elegans* strains. An alkaline phosphatase-labeled cDNA probe specific to Orsay virus RNA1 was used. Ribosomal RNA stained with methylene blue served as a loading control. Right panel: Quantification of Orsay virus RNA1 levels from the northern blot using ImageJ. B. Size distribution and 5′ nucleotide composition of Orsay virus RNA1-derived siRNAs in the same worm strains shown in panel A. C. Size distribution and 5′ nucleotide composition of Orsay virus RNA1-derived siRNAs following treatment with Terminator phosphatase to clone secondary vsiRNAs with 5′ tri-phosphorylate during sequencing library construction. Worm strains analyzed are the same as in panel A. D. Left panel: Quantification of 22–24 nt Orsay virus RNA1-derived primary siRNAs in the same worm strains as in panel A. Right panel: Quantification of secondary siRNAs derived from Orsay virus RNA1 in the same strains. Read counts were normalized to one million total miRNAs.

To quantify secondary vsiRNAs in wild-type N2 and RNAi-defective mutants, we performed subtraction analyses: the abundance of primary vsiRNAs detected in direct cloning libraries (which exclude secondary vsiRNAs) was subtracted from that detected in enzyme-treated libraries (which capture both primary and secondary vsiRNAs). Consistent with the size distribution data (Figure 2C), secondary vsiRNAs accumulated to the highest levels in *rsd-6*(*kd622*) mutants (Figure 2D, right panel). In contrast, they were barely detectable in *rde-1* mutants and were present at significantly reduced levels in *rrf-1* animals compared to wild-type N2 (Figure 2D, right panel). These results demonstrate that *rsd-6* is not required for secondary vsiRNA production.

### Viral replication is further enhanced in rrf-1;rsd-6 double mutants compared to rrf-1 single mutants

The observation that both primary and secondary vsiRNAs accumulate to high levels in *rsd-6* mutants suggests that RSD-6 may contribute to antiviral RNAi by facilitating the activity of secondary vsiRNAs. To test this, we compared OrV replication in *rde-1;rsd-6* and *rrf-1;rsd-6* double mutants with their respective single mutants. Because secondary vsiRNAs are still detectable, though at low levels, in *rrf-1* mutants but nearly absent in *rde-1* mutants, we reasoned that loss of *rsd-6* would have little effect in the *rde-1* background, but would abolish the antiviral activity of secondary vsiRNAs generated in *rrf-1* mutants. Accordingly, we predicted that OrV replication would be further enhanced in *rrf-1;rsd-6* double mutants compared with *rrf-1* single mutants, but unchanged in *rde-1;rsd-6* double mutants compared with *rde-1* single mutants.

Consistent with this hypothesis, OrV replication was markedly elevated in *rrf-1;rsd-6* double mutants compared with *rrf-1* single mutants, whereas no difference was observed between *rde-1;rsd-6* and *rde-1* mutants (Figure 3A). As expected, the high levels of viral replication in double mutants correlated with strong accumulation of primary vsiRNAs derived from OrV RNA1 (Figure 3B and 3D, left panel). Interestingly, while secondary vsiRNAs were barely detectable in *rde-1;rsd-6* double mutants, they were readily detected in *rrf-1;rsd-6* double mutants (Figure 3C and 3D, right panel). This indicates that secondary vsiRNAs can be generated independently of *rrf-1* during antiviral RNAi, and that their antiviral function requires RSD-6.

**Figure 3.**
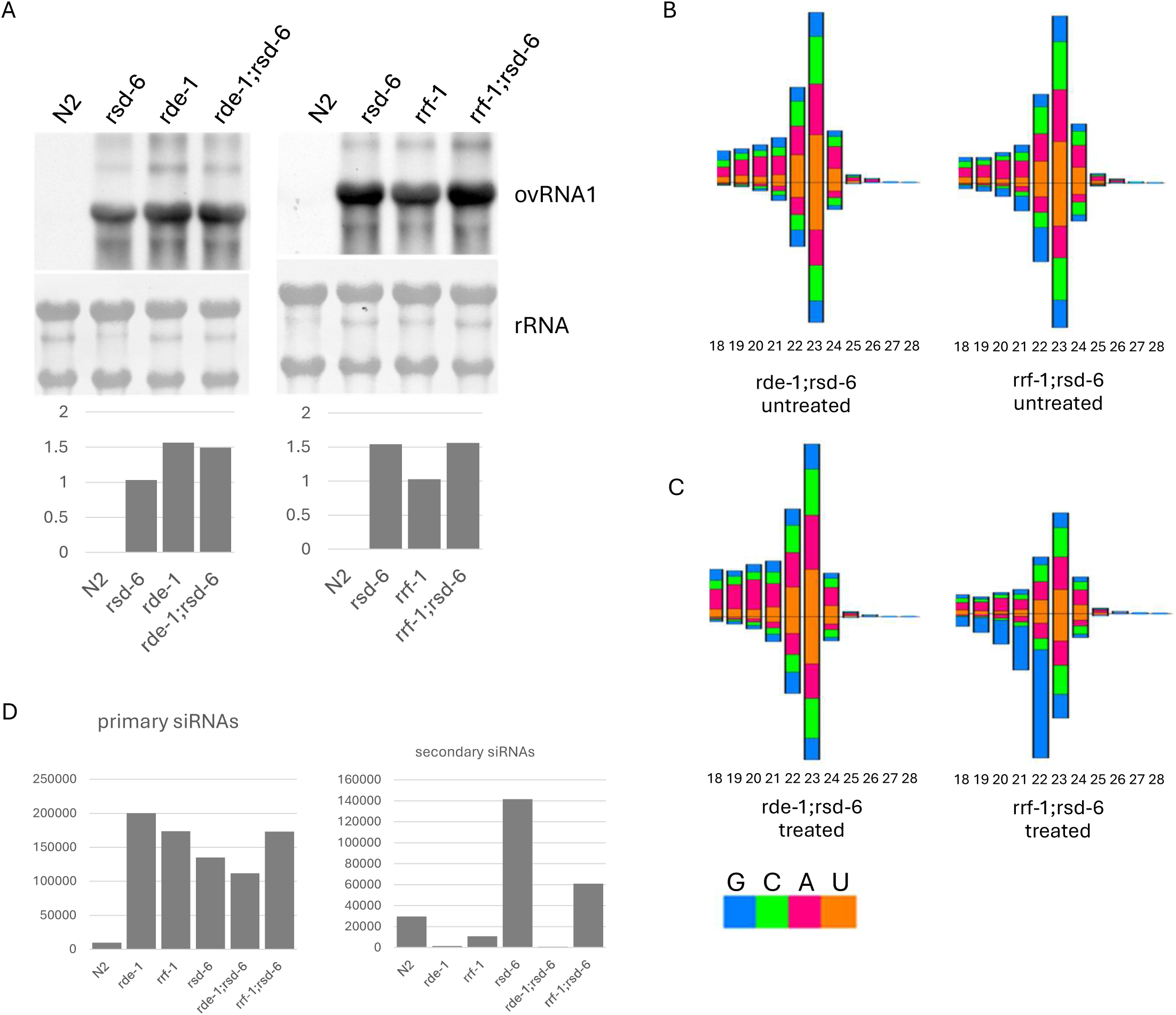
RSD-6 contributes to the function of virus-derived secondary siRNAs. A, Upper panel, northern blot detection of OrV genomic RNA1 in the indicated *C. elegans* strains. Experimental conditions were as described in Figure 2A. Lower panel, quantification of OrV RNA1 levels from the upper panel, analyzed using ImageJ. B, Size distribution and 5′ nucleotide composition of OrV RNA1-derived vsiRNAs in the indicated double mutants. Data represent direct sequencing of small RNAs (primary siRNAs). C, Size distribution and 5′ nucleotide composition of OrV RNA1-derived vsiRNAs in the same double mutants as in (A), but with Terminator phosphatase-treated small RNA samples to detect secondary siRNAs. D, Left panel, quantification of 22-24 nt OrV RNA1-derived primary vsiRNAs in the indicated strains. Read counts were normalized to one million total miRNA reads. Right panel, quantification of OrV RNA1-derived secondary vsiRNAs in the indicated strains. Secondary siRNA reads were similarly normalized to one million total miRNAs.

### The N-terminal tandem domains, but not the C-terminal Tudor domain, of RSD-6 are required for antiviral function

AlphaFold3 structure prediction suggests that *C. elegans* RSD-6 contains two tandem Tudor domains at C-terminus and three tandem domains (NTDs) of unknown function N-terminus (Figure S3). Each NTD consists of three α-helices and two β-sheets that form a triangular pyramid, with the β-sheets at the base. The functional contributions of these domains to antiviral RNAi remain unclear. To address this, we generated truncation mutants lacking specific domains (Figure 4A) and expressed them in *rsd-6(kd622)* mutants carrying the *FR1gfp* replicon transgene. Expression of these RSD-6 derivatives, driven by a heat-inducible promoter (as in Figure 1D), was verified by Western blot (Figure 4B). Antiviral activity was then assessed by measuring *FR1gfp* genomic and subgenomic RNA levels using RT-qPCR.

**Figure 4.**
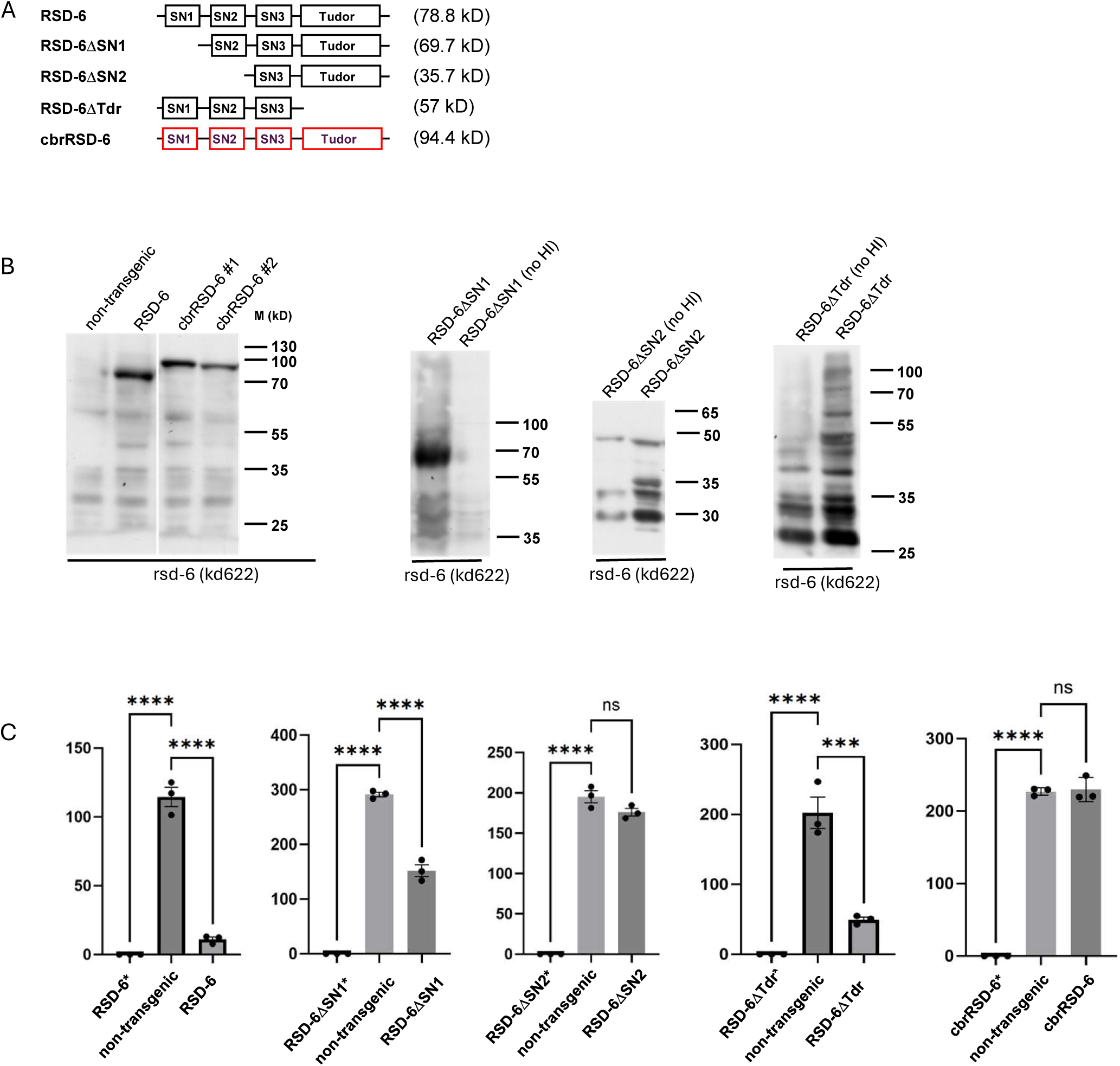
Functional domain characterization of RSD-6. (A) Schematic domain structure and predicted molecular weight (kDa) of RSD-6 and its derivatives. SN: staphylococcal nuclease domain; Tudor: Tudor domain; cbrRSD-6: C. briggsae RSD-6 ortholog. (B) Western blot analysis of HA-tagged RSD-6 and its derivatives as indicated. No HI, no heat induction; * indicates non-induced samples). (C) Quantitative analysis of FR1gfp replicon RNA1 accumulation by RT-qPCR in rsd-6(kd622) mutants expressing either wild-type RSD-6 or its derivatives. All strains contained both the FR1gfp replicon transgene and rsd-6(kd622) null allele. Statistical significance: ***, p < 0.0001, ****, p < 0.00001.

Overexpression of wild-type RSD-6 fully restored antiviral function in *rsd-6(kd622)* mutants (Figure 4C). In contrast, mutants lacking the first NTD, or both the first and second NTDs, showed progressively weaker antiviral activity, indicating that all three NTDs are required for full antiviral potency. By comparison, removal of the Tudor domains did not impair antiviral function, suggesting it is dispensable for antiviral defense. Interestingly, *C. briggsae* RSD-6 shares the same domain architecture as *C. elegans* RSD-6, but AlphaFold3 predicts a distinct spatial arrangement (Figure S3). To test for functional conservation, we expressed HA-tagged *C. briggsae* RSD-6 in the same system. Although expression was confirmed (Figure 4C), it failed to suppress *FR1gfp* replication, unlike *C. elegans* RSD-6. These findings suggest that species-specific differences in spatial domain organization underlie the divergent antiviral activities of RSD-6.

### RSD-6 is a key modulator of Orsay virus pathogenesis

In wild-type N2 animals, Orsay virus replicates at low levels and causes no detectable disease symptoms, likely due to the absence of RNAi suppression activity (7, 8, 16). Notably, neither the viral coat protein nor the C-terminal domain of the coat–delta fusion protein exhibits RNAi suppression when overexpressed in transgenic animals (14). However, in *drh-1; rde-1; rde-4* triple mutants, which are free of RNAi responses, Orsay virus replicates robustly, leading to significantly reduced brood sizes and partial lethality (25, 26). To determine whether *rsd-6* similarly influences viral pathogenesis, we performed survival assays in *rsd-6* (*kd622*) mutants using established methods (27). When fed OP50 bacteria, *rsd-6*, *rde-1*, and *rrf-1* mutants exhibited survival rates comparable to N2 controls (Figure 5A). However, following Orsay virus infection, ∼60% of *rsd-6* and *rde-1* mutants died by 6 days post-inoculation (dpi), whereas N2 animals showed no clear disease symptoms (Figure 6B). *rrf-1* mutants also displayed reduced survival, though less severe than *rsd-6* or *rde-1* mutants. Notably, some of infected *rde-1*, *rrf-1*, and *rsd-6* mutants were able to lay eggs before death, preventing population collapse (data not shown). These findings demonstrate that *rsd-6*, like *rde-1* and *rrf-1*, is a critical determinant of Orsay virus-induced pathogenesis.

**Figure 5.**
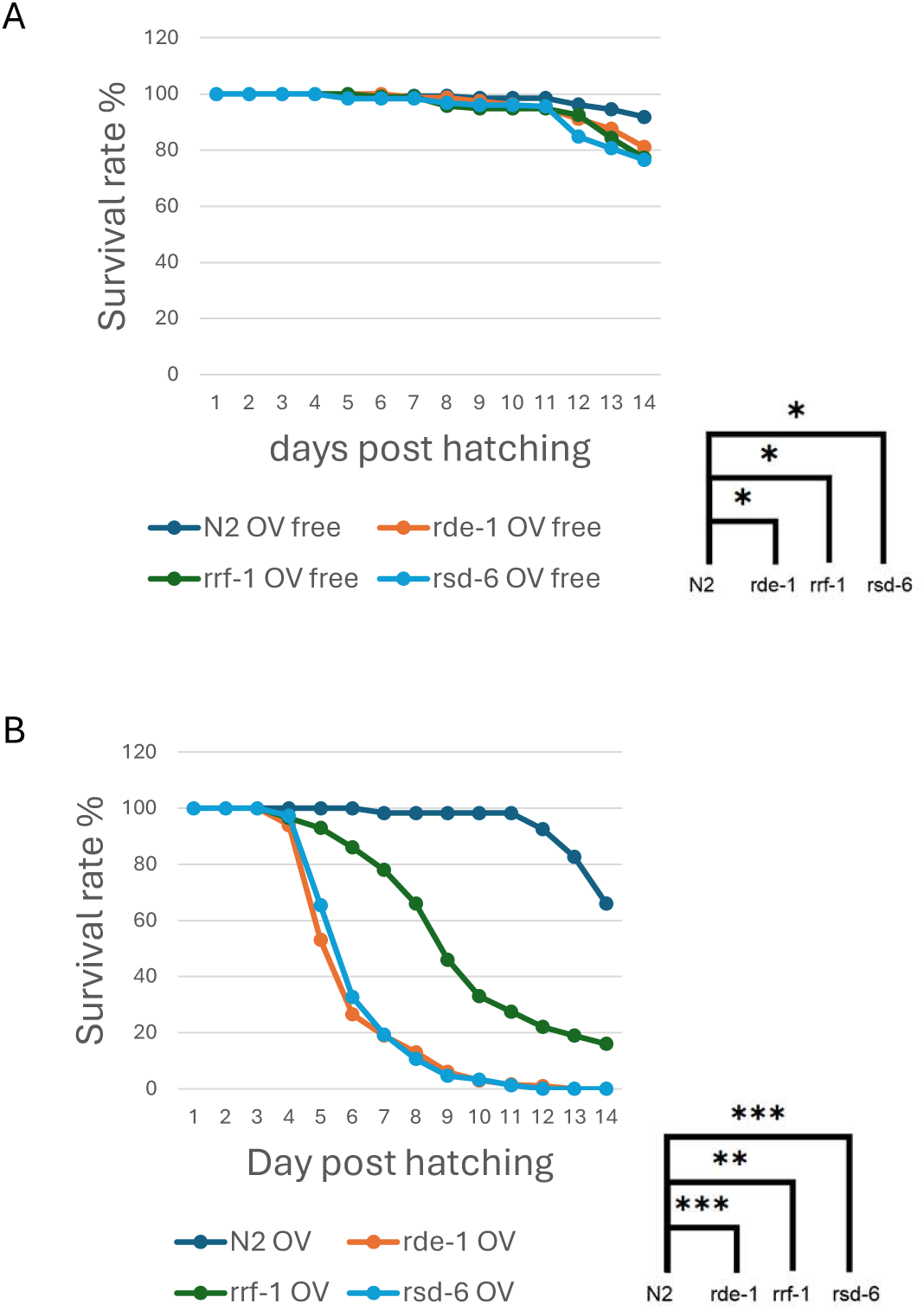
RSD-6 modulates viral pathogenesis in *C. elegans*. (A) Survival assay of mock-infected *C. elegans* strains. All worms were maintained on OP50 E. coli food. (B) Survival assay of Orsay virus-infected *C. elegans* strains as indicated. Animals were challenged with Orsay virus at the L1 stage, and survival rates were monitored every 24 hours post-infection. Statistical significance: *, *p* < 0.02; **, *p* < 0.002; ***, *p* < 0.0002; ****, *p* < 0.00002.

**Figure 6.**
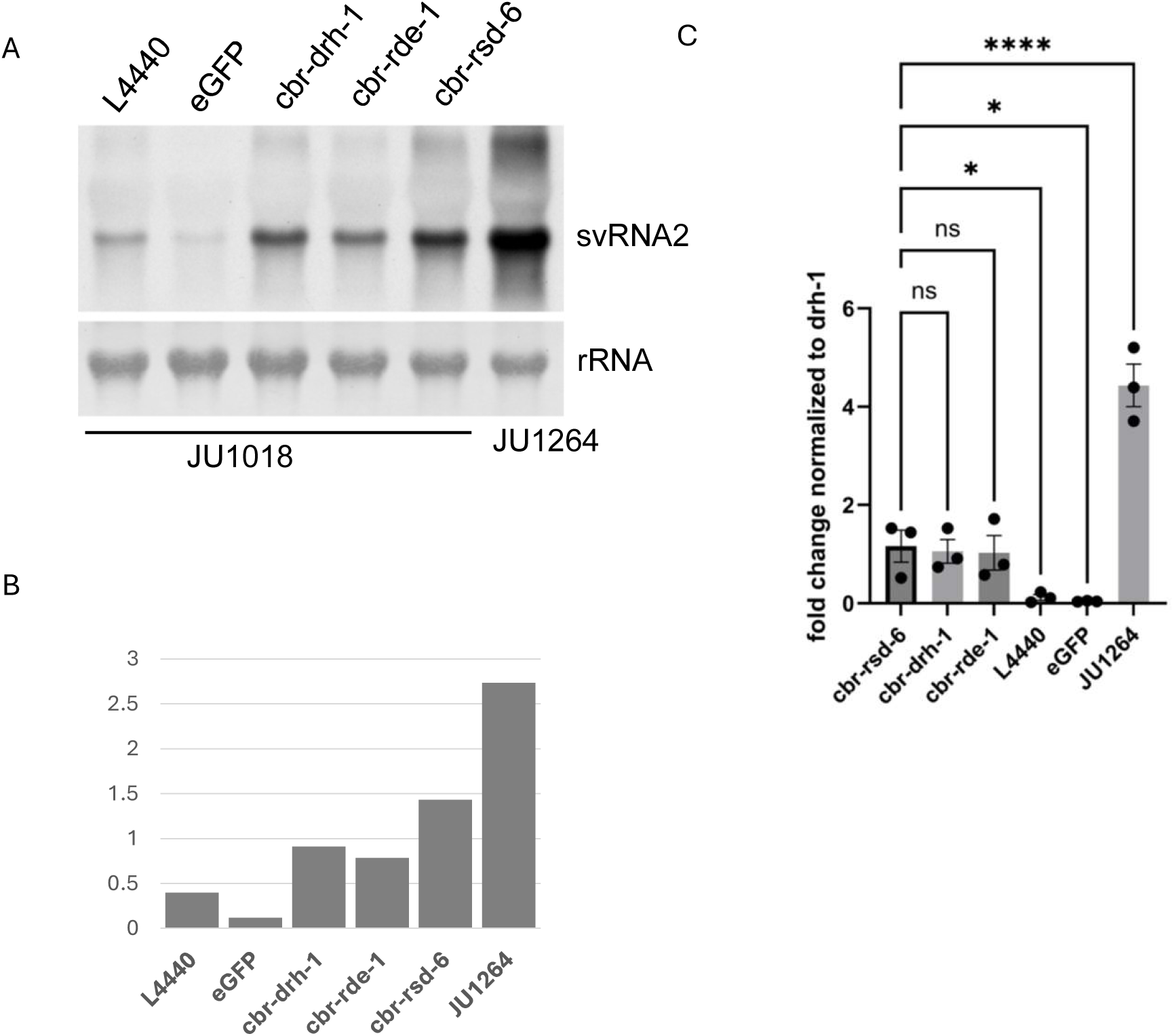
The *rsd-6* homolog contributes to antiviral immunity in *C. briggsae*. (A) Northern blot detection of Santeuil virus RNA2 in *C. briggsae* strains after RNAi knockdown of indicated genes. Total RNA (5 µg per lane) from Santeuil virus-infected animals was probed with cDNA derived from the viral RNA2 segment. JU1018, *C. briggsae* strain with *C. elegans sid-2* transgene. JU1264, original Santeuil virus host. L4440, empty vector; eGFP, dsRNA targeting eGFP. rRNA, methylene blue-stained ribosomal RNA serving as the loading control. svRNA2, Santeuil virus RNA2. B. Quantification of Santeuil virus RNA2 levels from (A) using ImageJ. (C) RT-qPCR quantification of Santeuil virus RNA2 in JU1018 worms fed with dsRNA targeting indicated genes. Data represent mean ± SD of three independent experiments. Statistical analysis: *, *p* < 0.05; *****, *p* < 0.00005; ns, not significant.

### RSD-6 homologue mediates antiviral defense in C. briggsae

The *rsd-6* gene is conserved across nematodes, suggesting an important functional role. Previous work showed that among the genes differentially expressed in response to Orsay virus infection in *C. elegans*, 58 are also differentially expressed in *C. briggsae* during SANTV infection (28), raising the possibility that the antiviral role of *rsd-6* is conserved across *Caenorhabditis* species. To test this, we silenced the *rsd-6* homolog in *C. briggsae* by feeding RNAi and monitored SANTV replication using northern blotting and RT-qPCR. Because *C. briggsae* is generally less sensitive to ingested dsRNA, we used the JU1018 strain, which carries the *C. elegans sid-2* transgene and exhibits enhanced feeding RNAi sensitivity, as confirmed by *dpy-11* silencing (data not shown) (29, 30). As shown in Figure 6A and B, feeding JU1018 worms with HT115 bacteria expressing dsRNA targeting *C. briggsae drh-1* or *rde-1* significantly enhanced SANTV replication. A similar increase was observed upon *rsd-6* silencing, whereas no effect was seen in controls treated with eGFP dsRNA or the empty L4440 vector. Consistent with these results, RT-qPCR quantification confirmed elevated viral RNA levels following *rsd-6* knockdown (Figure 6C). Together, these findings demonstrate that the antiviral function of *rsd-6* is evolutionarily conserved in *Caenorhabditis* nematodes.

## Discussion

*rsd-6* has been implicated in multiple siRNA-mediated RNA silencing pathways in *C. elegans*, including transposon control, antiviral defense, and gene silencing triggered by artificial dsRNAs (12, 31–34). However, its precise role within these pathways has remained unclear. Here, we show that virus-derived secondary siRNAs accumulate at high levels in *rsd-6* mutants, suggesting a role for RSD-6 downstream of secondary siRNA biogenesis. Consistent with this, secondary vsiRNAs produced in an *rrf-1*–independent manner remained detectable in *rrf-1;rsd-6* double mutants. Importantly, viral replication was further enhanced in *rrf-1;rsd-6* double mutants compared with *rrf-1* single mutants, indicating that RSD-6 is required for the activity of secondary vsiRNAs generated through both *rrf-1*–dependent and *rrf-1*–independent mechanisms. This conclusion is further supported by the observation that Orsay virus infection caused more severe disease symptoms in *rsd-6* mutants than in *rrf-1* mutants. Moreover, using a *C. briggsae* transgenic line with enhanced feeding RNAi activity, we found that downregulation of *C. briggsae rsd-6* expression compromised antiviral defense, demonstrating that the antiviral function of RSD-6 is conserved across *Caenorhabditis* nematodes.

By normalizing to the accumulation of miRNAs, whose biogenesis is known not to be affected by defects in antiviral RNAi pathway (23), we were able to evaluate the abundance of vsiRNAs in different genetic backgrounds. Notably, we found that Orsay virus RNA1 genome segment produced significantly fewer primary vsiRNAs but at the same time yielded more secondary vsiRNAs compared to the RNA2 genome segment (Figure S1). Given the relatively small size difference between these two genome segments, 3421 nucleotides vs 2574 nucleotides, we believe that an intrinsic mechanism yet to be identified may be in place to control the dynamics of vsiRNA production. It was also clear that, although their abundance varied significantly, primary vsiRNAs derived from each genome segment exhibit a unique pattern of genome coverage which remains unchanged in different genetic backgrounds (Figure S2). Viral genomic RNAs are replicated in replication organelles formed by viral replicases and cellular membrane (35). These genome replication organelles as physical barriers keep viral dsRNAs isolated and may protect them from being processed by Dicer and co-factors. Mature viral genomic RNAs carry packaging signal (36), which allows them to be specifically packaged into capsids and stay safe from being targeted by primary vsiRNA-loaded RDE-1. These observations suggest that physical association with viral and cellular products may protect viral single-stranded or double-stranded RNAs from being cleavage by RDE-8 or Dicer to different extents, thereby giving rise to the variations in vsiRNAs biogenesis. The unique pattern of genome coverage of primary vsiRNAs may reflect such a protection mechanism in which only unprotected viral dsRNAs are processed into primary vsiRNAs and the abundance of primary vsiRNAs correlate proportionally to the length of unprotected viral dsRNAs. The RNA2 genome segments of Orsay virus encodes the structure proteins which need to be produced at significantly higher levels compared to the replicase encoded by RNA1. This means that the RNA2 segment needs to get engaged in translation at a much higher frequence compared to RNA1 segment and the physical association with translation machinery may protect them from being targeted for cleavage by RDE-8. This may explain why much less secondary vsiRNAs are derived from RNA2. RNA-seq aiming to detect viral RNAs with poly(UG) tails should help address this hypothesis.

By normalizing vsiRNA levels to miRNAs, whose biogenesis is unaffected by defects in the antiviral RNAi pathway (23), we assessed vsiRNA abundance across different genetic backgrounds. Interestingly, the Orsay virus RNA1 genome segment produced significantly fewer primary vsiRNAs but generated more secondary vsiRNAs compared to the RNA2 segment (Figure S1). Given the modest size difference between these two genome segments (3421 vs. 2574 nucleotides), this disparity likely reflects an intrinsic mechanism regulating vsiRNA production. Moreover, although their abundance varied, primary vsiRNAs from each genome segment exhibited distinctive and reproducible coverage patterns that remained unchanged across genetic backgrounds (Figure S2).

Orsay virus genomic RNAs are replicated within replication organelles formed by viral replicases and host membranes (35). These organelles may act as physical barriers, shielding viral dsRNAs from processing by Dicer and its cofactors. In addition, mature genomic RNAs carry packaging signals (36), enabling them to be selectively encapsidated and protected from cleavage by RDE-8 guided by primary vsiRNAs. Together, these physical associations with viral and host factors may differentially shield viral ssRNAs and dsRNAs from processing by RDE-8 or Dicer, thereby generating the observed variation in vsiRNA biogenesis. The distinct coverage patterns of primary vsiRNAs likely reflect such protection, with only unshielded viral dsRNAs accessible for Dicer cleavage. Accordingly, the abundance of primary vsiRNAs may be proportional to the length of exposed dsRNA regions. Notably, the RNA2 segment encodes structural proteins that must be produced at much higher levels than the RNA1-encoded replicase. Consequently, RNA2 is engaged in translation machineries more frequently, and association with the translation machinery may protect it from RDE-8 mediated cleavage. This provides a plausible explanation for why RNA2 yields fewer secondary vsiRNAs. Future RNA-seq analyses designed to detect viral RNAs carrying poly(UG) tails should help test this hypothesis.

RSD-6 was originally identified as a factor required for systemic RNAi in *C. elegans* (34). However, no antiviral function of systemic RNAi has been demonstrated to date, suggesting that RSD-6 may contribute to systemic RNAi and antiviral silencing through distinct mechanisms. RSD-6, together with RSD-2, has also been shown to be essential for the biogenesis of endogenous 22G RNAs (37). By maintaining a pool of these RNAs, RSD-6 and RSD-2 promote germ cell immortality, particularly under stress conditions (31). In *rsd-6* and *rsd-2* mutants, 22G RNAs targeting spermatogenesis genes are perturbed, and silencing of transposons and other repetitive loci is compromised, implicating both proteins in nuclear gene silencing within the germline.

Although secondary vsiRNAs share the same size and terminal structure as endogenous 22G RNAs, their biogenesis is mechanistically distinct. For example, key genes required for endogenous 22G RNA production, such as *ergo-1* and *rrf-3*, are dispensable for antiviral RNAi (38). Moreover, FHV and Orsay virus, both positive-strand RNA viruses, replicate in the cytoplasm of somatic cells, where secondary vsiRNAs are generated and act. Therefore, the distinct roles of RSD-6 in secondary vsiRNAs and endogenous 22G RNAs production likely reflect the different mechanisms RSD-6 deploys to contribute to nuclear gene silencing and cytoplasmic antiviral RNAi.

RSD-6 contains three tandem N-terminal domains (NTDs) of unknown function and two tandem Tudor domains at the C-terminus. How these domains contribute to antiviral function remains unclear. To address this, we performed functional rescue assays using RSD-6 truncation mutants, in which restoration of antiviral activity was assessed upon overexpression of each mutant construct. In this assay, removal of one or two NTDs led to a progressive loss of antiviral activity (Figure 4), indicating that these domains are critical for mediating antiviral defense. Surprisingly, deletion of the C-terminal Tudor domains did not compromise rescue activity, suggesting that the Tudor domains are dispensable under these conditions.

Tudor domain proteins typically act as molecular adaptors, promoting protein–protein interactions and facilitating the assembly of macromolecular complexes through recognition of methylated arginine or lysine residues (39). For example, in the endogenous RNAi pathway, the tandem Tudor protein ERI-5 tethers the RdRP RRF-3 to DCR-1, thereby potentiating siRNA biogenesis (40). By analogy, the Tudor domains of RSD-6 may cooperate with the NTDs to mediate protein–protein interactions required for RISC assembly with secondary siRNAs. However, this function may become dispensable when RSD-6ΔTudor is overexpressed, presumably high protein abundance could have compensated for the loss of Tudor-mediated interactions.

Plant genomes encode multiple Dicer-like proteins, each responsible for producing siRNAs of specific sizes (41, 42). By contrast, *C. elegans* encodes only a single Dicer, yet primary vsiRNAs of different size classes are consistently detected (9, 14). Which of these vsiRNAs play the dominant role in directing viral RNA cleavage remains unclear. The inability of the P19 viral suppressor, which binds 21-nt siRNAs, to inhibit antiviral RNAi in *C. elegans* suggests that vsiRNAs longer than 21 nt are the major contributors to antiviral silencing (14). Based on this, we focused our quantitative analyses on vsiRNAs ranging from 22 to 24 nt, which are produced with the highest abundances.

Consistent with previous studies, abundant primary vsiRNAs were detected in *rde-1* and *rrf-1* mutants, both of which are dispensable for primary vsiRNA production (8, 9) (Figure 2B, D). Similarly, *rsd-6* mutants accumulated primary vsiRNAs at comparable levels and with similar size distributions and genome coverage, indicating that RSD-6 acts downstream of primary vsiRNA biogenesis. By contrast, the analysis of secondary vsiRNAs revealed distinct patterns. As expected, secondary vsiRNAs were nearly absent in *rde-1* mutants, detectable at low levels in *rrf-1* mutants, and accumulated to high levels in *rsd-6* mutants (Figure 2C, D). These findings indicate that RSD-6 functions downstream of secondary vsiRNA biogenesis. Consistently, while secondary vsiRNAs were barely detectable in *rde-1;rsd-6* double mutants, they were readily detected in *rrf-1;rsd-6* double mutants, confirming that RSD-6 is not required for the production of secondary vsiRNAs generated via either *rrf-1*–dependent or –independent pathways.

*C. briggsae* RSD-6 shares the same overall domain organization as *C. elegans* RSD-6 and is also required for antiviral defense (Figure 6). However, overexpression of *C. briggsae* RSD-6 failed to restore antiviral defense in *C. elegans rsd-6(kd622)* null mutants. Notably, *C. briggsae* RSD-6 is larger (813 vs. 689 amino acids) and contains longer linker sequences between its N-terminal tandem domains. Structural predictions using AlphaFold3 suggest that, in *C. elegans* RSD-6, the N-terminal tandem domains adopt a distinct spatial orientation relative to the C-terminal tandem Tudor domains (Figure S3). This raises the possibility that species-specific differences in the spatial arrangement of these domains determine functional specificity, perhaps by enabling interactions with different co-factors critical for antiviral defense in each nematode species. Given that the tandem Tudor domains of *C. elegans* RSD-6 are dispensable for antiviral activity (Figure 4), it will be interesting to test whether antiviral defense in *rsd-6* null mutants can be restored by a chimeric protein combining the N-terminal tandem domains of *C. briggsae* RSD-6 with the linker sequences of *C. elegans* RSD-6.

## Materials and Methods

### Animal genetics and maintenance

The *C. elegans* Bristol N2 isolate served as the reference wild-type strain in this study. All strains were maintained at room temperature (22°C) on standard Nematode Growth Medium (NGM) seeded with OP50 *E. coli* as a food source. The mutant strains used in this study, including *drh-1(ok3495)*, *rde-1(ok569)*, rde-3 (ne3370), *rde-4(yt4350),*and *rrf-1 (ok589) w*ere either obtained from the Caenorhabditis Genetics Center (CGC) or generated in our laboratory (See table S2 for details). The genotypes for strains used in this study were verified by PCR followed by Sanger sequencing. Primers used for genotyping are listed in Table S3.

### Genetic crosses and genetic complementation

Males were obtained either through natural occurrence or by treating hermaphrodites with 10% ethanol (room temperature, 15 minutes) to suppress heat-inducible viral replicon expression in transgenic strains. All crosses were performed on NGM plates supplemented with 100 µg/ml ampicillin. The male-to-hermaphrodite ratio was adjusted based on the transgene or genetic allele involved. Homozygous F2 progeny were identified through PCR genotyping and reconfirmed using Sanger sequencing.

### Feeding RNAi

The feeding RNAi assay was carried out using *C. briggsae* strain JU1018. JU1018 carries a transgene derived from *C. elegans sid-2* which makes JU1018 more sensitive to feeding RNAi. For eGFP feeding RNAi the full-length coding sequence of eGFP was inserted into L4440 and the resulting construct was transferred into HT115 bacteria which will produce eGFP dsRNA upon IPTG induction. 4 hours of IPTG treatment was used to induce the dsRNA production before the medium being seeded on the standard ampicillin NGM agar plates. Gravid hermaphrodites were placed on feeding RNAi plates to lay eggs at the room temperature (RT). Total RNA samples were then prepared for fed F1 progenies after they reach to young or fully developed adults. As negative control, worms were provided with feeding RNAi bacteria containing empty L4440 vector.

### Function rescue

To create constructs for *rsd-6* function rescue assay, coding sequence for *rsd-6* or its derivatives was amplified using total cDNA templates prepared for wildtype N2 or *C. briggsae* reference strain JU1085. The amplified coding sequences were cloned into an overexpression vector under the control of *hsp-16.41* heat inducible promoter through Gibson assembly. The rescue constructs were injected into *rsd-6* (*kd622*) mutants at 10 ng/µl together with a co-injection marker, Pmyo-2::mCherry (20ng/ µl) and 1 kb Plus DNA Ladder (New England Biolabs, N3200L) (170 ng/ µl). Transgenic lines with a transmission rate of 30-50% were used for viral replication tests.

### CRISPR/Cas9 editing of C. elegans animals

*rsd-6* null allele kd622 was generated using CRISPR-Cas9 genome editing technique following the protocol as described previously (43). Two guide RNAs (crRNAs), were used to specify left and right cut sites in the rsd-6 locus so that the fragment in between will be removed. crRNAs were designed using the online tool provided by Integrated DNA Technologies (IDT). ssDNA oligo donor designed for homology directed repair (HDR) consisted of 35 nucleotides upstream of left cut site annealed to 35 nucleotides downstream of right cut site. Two crRNAs (rsd6gRNA1 and rsd6gRNA2) were designed for targeting the rsd-6 locus. A ssDNA oligo donor was designed and used as repair template (rsd-6 ssODN). The injection mix was prepared by combining the Cas9 protein, tracrRNA, rsd-6 crRNAs, repair template and PRF4 plasmid. The progeny of injected animals exhibiting roller phenotype were singled out and used for PCR-verification of CRISPR/Cas9-editing at the targeted locus.

### Structure prediction of RSD-6 from C. elegans or C. briggsae

Structures of wildtype and mutant VIRO-9 were predicted using AlphaFold2 (44). The output PDB files were visualized and annotated using UCSF Chimera (45). Predicted structures of the wildtype RSD-6 proteins from *C. elegans* and *C. briggsae* were colored in green and beige respectively. Overlay of the predicted structures were created in UCSF Chimera.

### FR1gfp replication induction and Orsay virus infection

At least three independent experiments were performed for northern blotting assays to detect FR1gfp or Orsay virus replication. Induction of FR1gfp replication in transgenic animals was achieved by heat-shocking synchronized young adult animals at 33°C for 1 hour and 45 minutes, followed by incubation at 25°C for up to 48 hours. Green fluorescence production was visualized and recorded using a dissecting microscope with UV illumination. Orsay virus was maintained in *rsd-2* mutants carrying a GFP transgene under the control of the *pals-5* promoter, which induces GFP expression upon Orsay virus infection (46). To collect viral particles, infected animals were washed from NGM plates using 5 mL autoclaved ddH₂O per plate. The suspension was centrifuged at 10,000 × g to separate viral particles from animals and bacterial cells, and the resulting supernatant was filtered through a 0.22 μm filter unit. The filtrate could be directly mixed with OP50 food for virus inoculation or stored in 20% glycerol at −80°C for future use. Santeuil virus was maintained using *C. briggsae* JU1264 strain. Virus filtrate preparation and infection assay was performed as described above.

### RNA preparation and viral RNA detection

Detection of FR1gfp and Orsay virus genomic and subgenomic RNAs was performed using previously described protocols (8). For viral RNA transcript detection via RT-qPCR, total high-molecular-weight RNA samples were used as input for cDNA synthesis with random primers, following the manufacturer’s instructions. The resulting cDNA was then subjected to qPCR amplification using virus-specific primers. Primers FR1gfp.F and FR1gfp.R were used for FR1gfp genomic RNA detection through RT-qPCR. Primers FR1gfp.subF and FR1gfp.subR were used for FR1gfp subgenomic RNA detection. The primers used for Orsay virus genomic RNA1 detection are OrV.FF and OrV.RR. Amplification of *act-1* served as internal control for all RT-qPCR assays. The primers used are act-1.Fa and act-1.R.

### Preparation and analysis of small RNA libraries

Infection of N2 and mutant worms with OrV was done as described previously (14, 25). For each sample, two separate small RNA libraries were prepared. To prepare a small RNA library capturing just the primary vsiRNAs, gel purified small RNA samples were directly used as inputs. To capture both primary and secondary vsiRNAs, gel purified small RNA was treated with RNA polyphosphatase (8µl RNA + 0.5µl RNA Polyphosphatase +0.5µl water + 1µl 10X RNA polyphosphatase buffer) for 30 minutes at 37°C. Small RNA libraries were prepared using NEBNext® Multiplex Small RNA Library Prep Set for Illumina following instructions from the manufacturer. Prepared small RNA libraries were quality tested using Agilent Bioanalyzer and sequenced on a NGS sequencing platform (Novaseq, Illumina). The sequencing data was analyzed using an in house developed pipeline on a Linux platform. All miRNAs were aligned to *C. elegans* miRNAs obtained from WormBase (WS240) and used to normalize vsiRNA reads. All virus small RNA reads were aligned to the OrV genome (GenBank identifiers [IDs] HM030970.2 and HM030971.2), allowing zero mismatches. Both sense and antisense vsiRNAs were normalized to one million of total miRNAs.

### Survival assay

Synchronized gravid adults were transferred to NGM plates seeded with *E. coli* OP50, either with or without Orsay virus. Worms were allowed to lay eggs overnight at 20°C, after which the adults were removed. F1 progeny were counted and maintained using the same food. Beginning the next day, surviving worms were transferred daily to fresh NGM plates to prevent interference from progeny. Worms were monitored daily for survival under a dissecting microscope, and individuals were scored as dead if they failed to respond to gentle touch with a platinum wire. The assay was conducted for 14 days, with worms maintained at 20°C throughout. Survival was expressed as a percentage of the initial population and plotted over time to generate survival curves. This assay was designed to evaluate the impact of Orsay virus on lifespan and overall survival of *C. elegans* under controlled laboratory conditions.

### Imaging microscopy

Both the red and green fluorescence images were recorded under the same exposure for each set of images. A Nikon digital camera Z6 II mounted on a Nikon SMZ1500 microscope was used to record all of the images.

## Acknowledgment

The authors thank the Caenorhabditis Genetics Center for providing some of the worm strains used in this study. We also extend our gratitude to undergraduate students Mary Hutson and Gabriel Loh for their assistance in preparing experimental materials and maintaining worm strains.

This work was supported by the National Institutes of Health (1R01GM119012-01A1 and 1R03AI171860-01A1).

---

**Figure S1.**
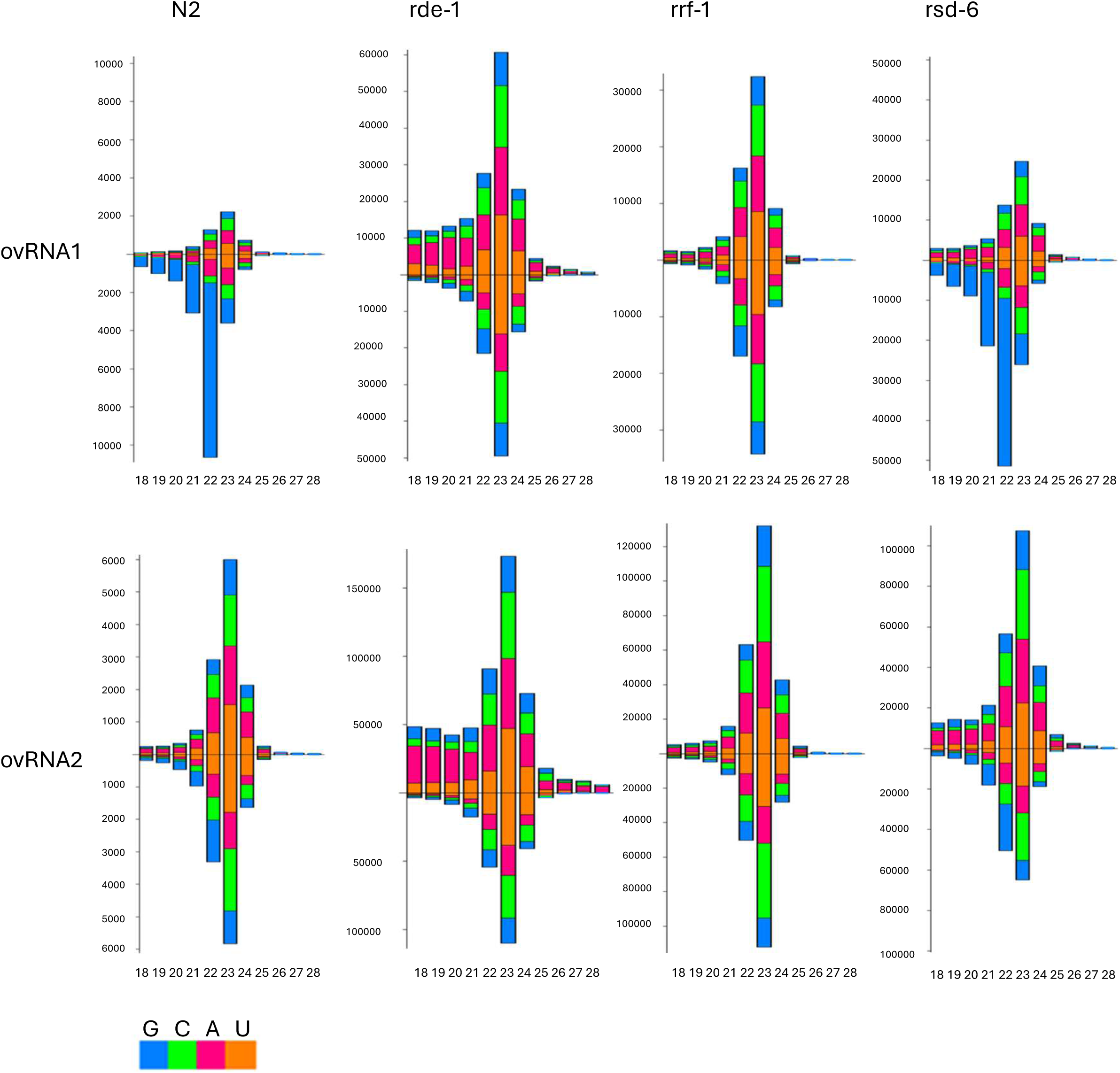
Distinct abundance and composition of siRNAs derived from Orsay virus RNA1 and RNA2. Shown are the size distribution and abundance of primary and secondary vsiRNAs mapped to Orsay virus RNA1 (ovRNA1) and RNA2 (ovRNA2) genomic segments. Read counts for siRNAs are normalized to one million total miRNA reads.

**Figure S2.**
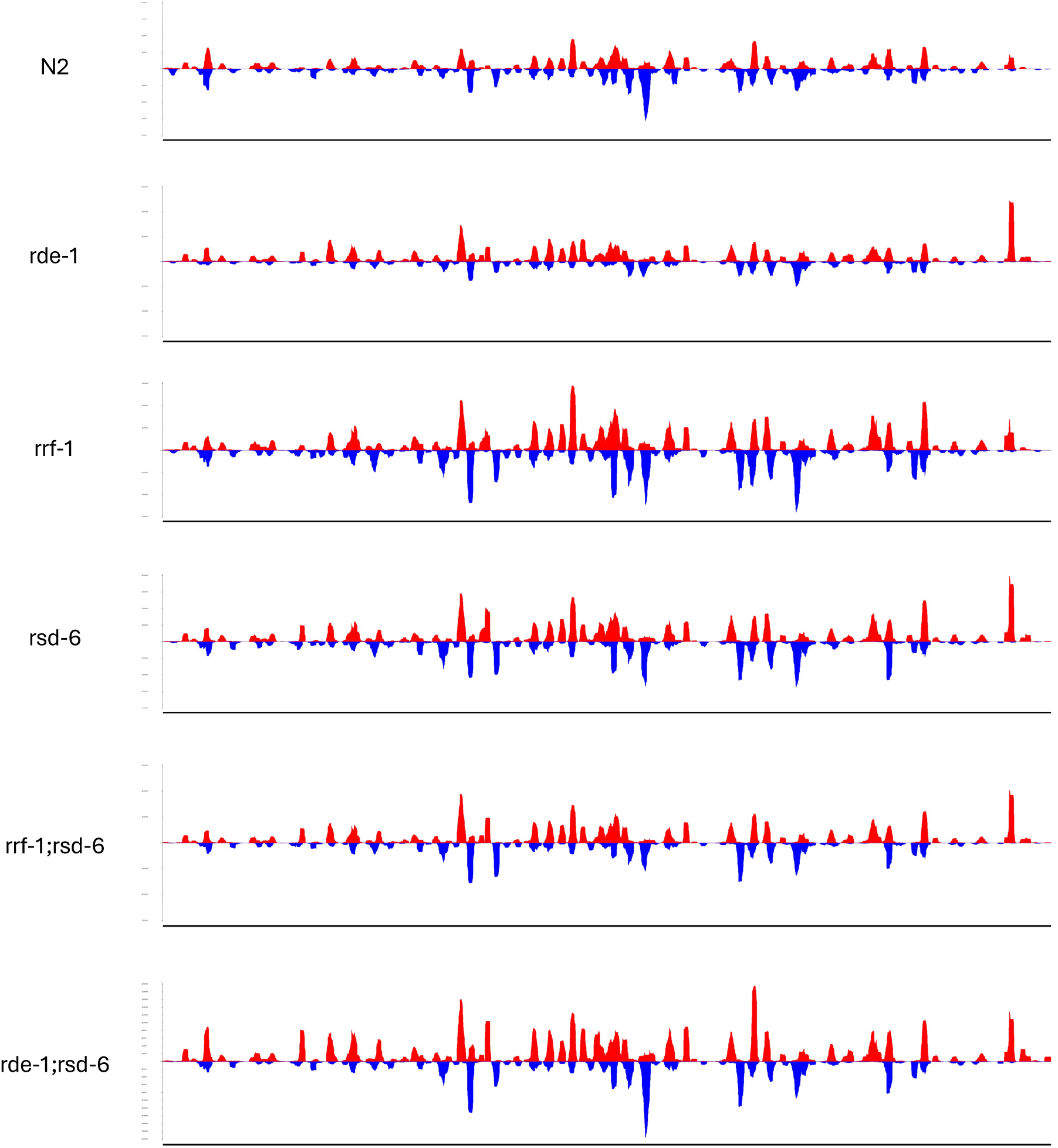
siRNAs mapped to OrV RNA1 exhibit the same pattern of distribution among N2 and mutant animals as indicated. Shown here is the mapping of 22-24 nt sense (red) and antisense (blue) vsiRNAs to the full-length OrV RNA1. The sequencing libraries were constructed using 5’ P-dependent protocol. The relative abundance was normalized to one million total miRNAs.

**Figure S3.**
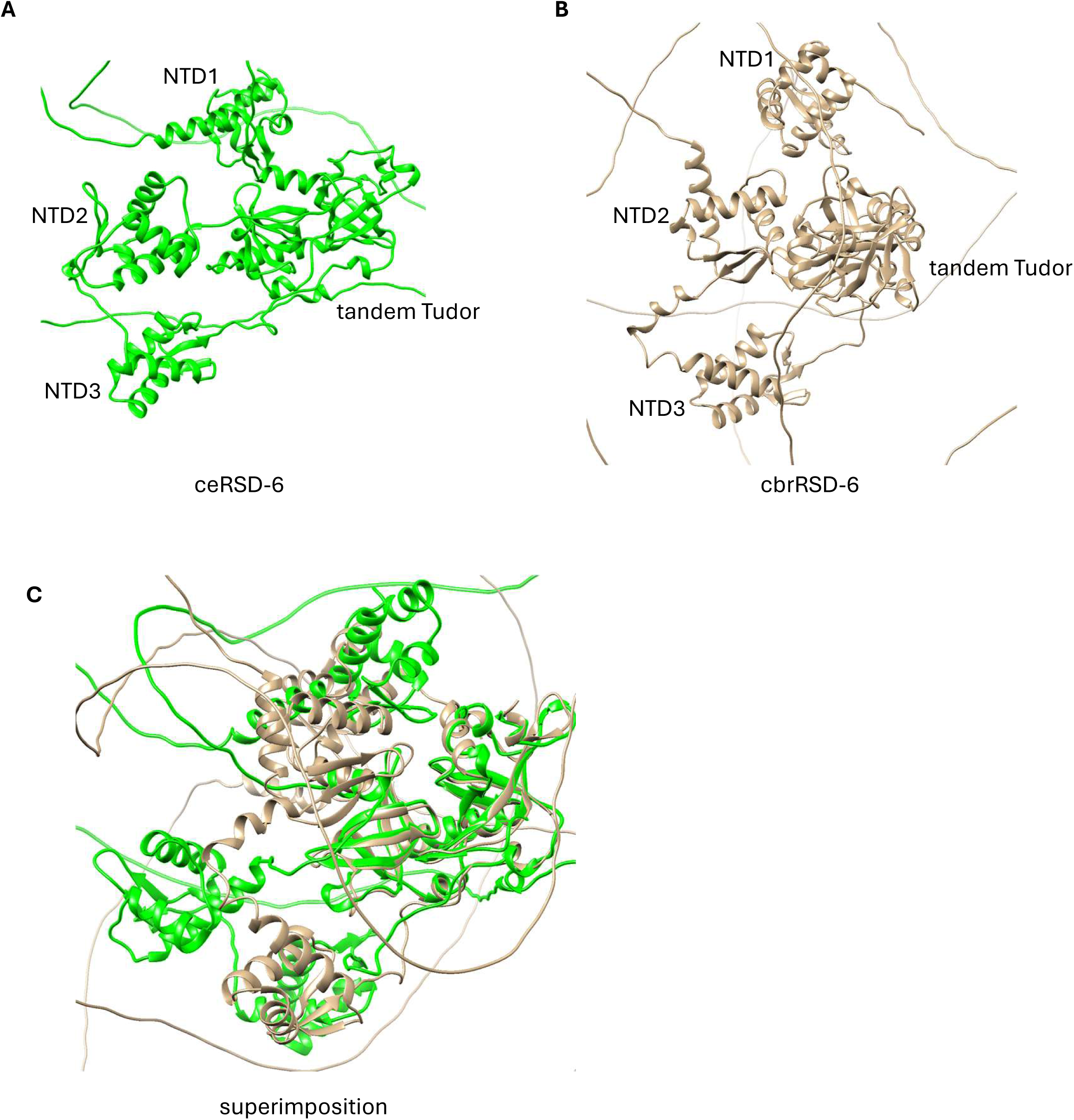
*C. elegans* RSD-6 and *C. briggsae* RSD-6 exhibit distinct function domain arrangements. A. AlphaFold3-predicted structures of wildtype *C. elegans* RSD-6 viewed in UCSF Chimera. C. AlphaFold3-predicted structures of wildtype *C. briggsae* RSD-6. C. Wildtype *C. elegans* RSD-6 superimposed on wildtype *C. briggsae* RSD-6 with the Tudor domain aligned in UCSF Chimera.

